# Identification of chemicals targeting dementia genes and pathways in the Comparative Toxicogenomics Database

**DOI:** 10.64898/2025.12.19.695572

**Authors:** Scarlet Cockell, Sean M. Harris, Rachel K. Morgan, Gary J. Patti, Erin B. Ware, Kelly M. Bakulski

**Affiliations:** Department of Epidemiology, School of Public Health, University of Michigan, Ann Arbor, MI; Department of Environmental Health Sciences, School of Public Health, University of Michigan, Ann Arbor, MI; Departments of Chemistry, Genetics and Medicine, Washington University School of Medicine, St. Louis, MO; Survey Research Center, Institute for Social Research, University of Michigan, Ann Arbor, MI

**Author notes:** **Corresponding author:** Kelly M. Bakulski, 1415 Washington Heights, Ann Arbor, MI, 48103.

**Keywords:** Dementia, Alzheimer’s disease, exposome, neurotoxicology, data-mining, chemical exposures

## Abstract

**Introduction:** Dementia is a public health challenge and exposures likely contribute to risk, though many have not been evaluated. We screened chemicals for enrichment with dementia genes and related pathways.

**Methods:** We obtained gene lists from the Comparative Toxicogenomics Database for 1,008 chemicals and nine dementia-related pathways (e.g., Alzheimer’s disease, tauopathies). We tested pairwise chemical-dementia gene enrichment using Fisher’s exact tests and proportional reporting ratios (PRR), accounting for multiple comparisons with false discovery rate (FDR<1×10^-6^).

**Results:** Of the chemicals tested, 742 (73.6%) were enriched for at least one dementia pathway and 15 with all nine pathways, including benzo(a)pyrene, ethanol, paraquat, and particulate matter. We observed 295 chemicals enriched for Alzheimer’s disease, including sodium arsenite (PRR = 57.9) and 305 enriched for tauopathies, including bisphenol A (PRR=37.7).

**Discussion:** We identified chemicals enriched for dementia pathways, suggesting broad classes of chemicals contribute to dementia.

## 1. Introduction

Dementia, characterized by progressive loss of neurons with accompanying deterioration of cognitive functions related to memory, language and behavior, is a significant public health challenge.^1,2^ In the United States alone, the cost of treating individuals with these disorders was $321 billion in 2022, and is predicted to rise above $1 trillion by 2050.^3,4^ Dementia subtypes include Alzheimer’s disease and frontotemporal lobe dementia, and the majority of persons living with dementia have multiple pathologies, termed mixed dementia.^4^ Important hallmarks of dementia include pathological protein aggregation, neuroinflammation, and aberrant proteostasis.^5^ In addition, emerging evidence suggests cellular stress, including endoplasmic reticulum (ER) stress and prolonged activation of the Integrated Stress Response pathway, play important roles in the etiology of dementia.^6,7^ Identifying factors contributing to shared dementia pathologies and to distinct dementia subtypes is critical for prevention.

While genetic factors play a role and risk loci for dementia subtypes including Alzheimer’s disease have been identified,^8^ increasing evidence points to environmental factors in the etiology of dementia.^9^ Components of air pollution, including particulate matter less than 2.5 microns in diameter (PM2.5) and nitric oxide, are associated with poorer executive function and memory.^10^ Pesticide exposure has also been linked to dementia risk in human cohorts,^11^ and support from animal and cell models demonstrates common forms including chlorpyrifos and dichlorodiphenyltrichloroethane (DTT) increase amyloid-β protein levels.^12,13^ Elevated exposures to metals (e.g., lead, cadmium, and manganese) are associated with cognitive decline in human epidemiology studies and elevated levels of molecular hallmarks such as aggregates of amyloid-β protein and tau neurofibrillary tangles in the brain, ER stress, oxidative stress, and neuroinflammation in animal models.^14^ Some exposures, such as intake of leafy greens, adherence to a Brain-healthy diet, and adequate glycemic control, are likely protective against dementia.^15^ Identifying additional environmental toxicants and dietary factors that contribute to disease risk or protection, as well as characterization of their cellular and molecular mechanisms involved in dementia development could reveal important therapeutic targets and new strategies for public health interventions.

Humans are exposed to a diverse array of chemicals throughout life, and only a small proportion of these have been thoroughly assessed for neurotoxic effects. The “exposome” refers to the totality of environmental exposures, including physical, biological, chemical and social factors, that an individual encounters throughout life.^16^ The chemical component of the exposome is vast and complex. As of January 30, 2025, the Environmental Protection Agency’s (EPA’s) Toxic Substances Control Act (TSCA) Inventory lists 86,741 chemicals, with 42,293 of these in active use in the United States as of 2025.^17^ Globally, more than 350,000 chemicals and chemical mixtures have been registered for use and production.^18^ It is not feasible to thoroughly screen such high numbers of chemicals using traditional laboratory experimental toxicology methods. Alternative approaches, such as data-mining and bioinformatic analysis of publicly available data, can identify chemicals and molecular pathways for targeted further study. Additionally, such approaches can provide important information for designing and interpreting epidemiology studies on environmental contributions to dementia.

The Comparative Toxicogenomics Database (CTD) provides expert curation of millions of interactions between chemicals, genes, and diseases reported in the scientific literature.^19^ The CTD offers a unique opportunity to assess a broad array of chemicals (15,167 chemicals contained in the CTD as of August 6, 2025) for known interactions with genes relevant for dementia, many of which are part of the human exposome.^16,19^ In this study, we identified chemicals with documented associations with genes that overlap with dementia risk and the associated molecular/cellular pathways (**Figure 1**). We expect these findings will provide a list of confirmed and novel chemicals to prioritize for future toxicology and epidemiology studies aimed at understanding environmental impacts on dementia.

**Figure 1.**
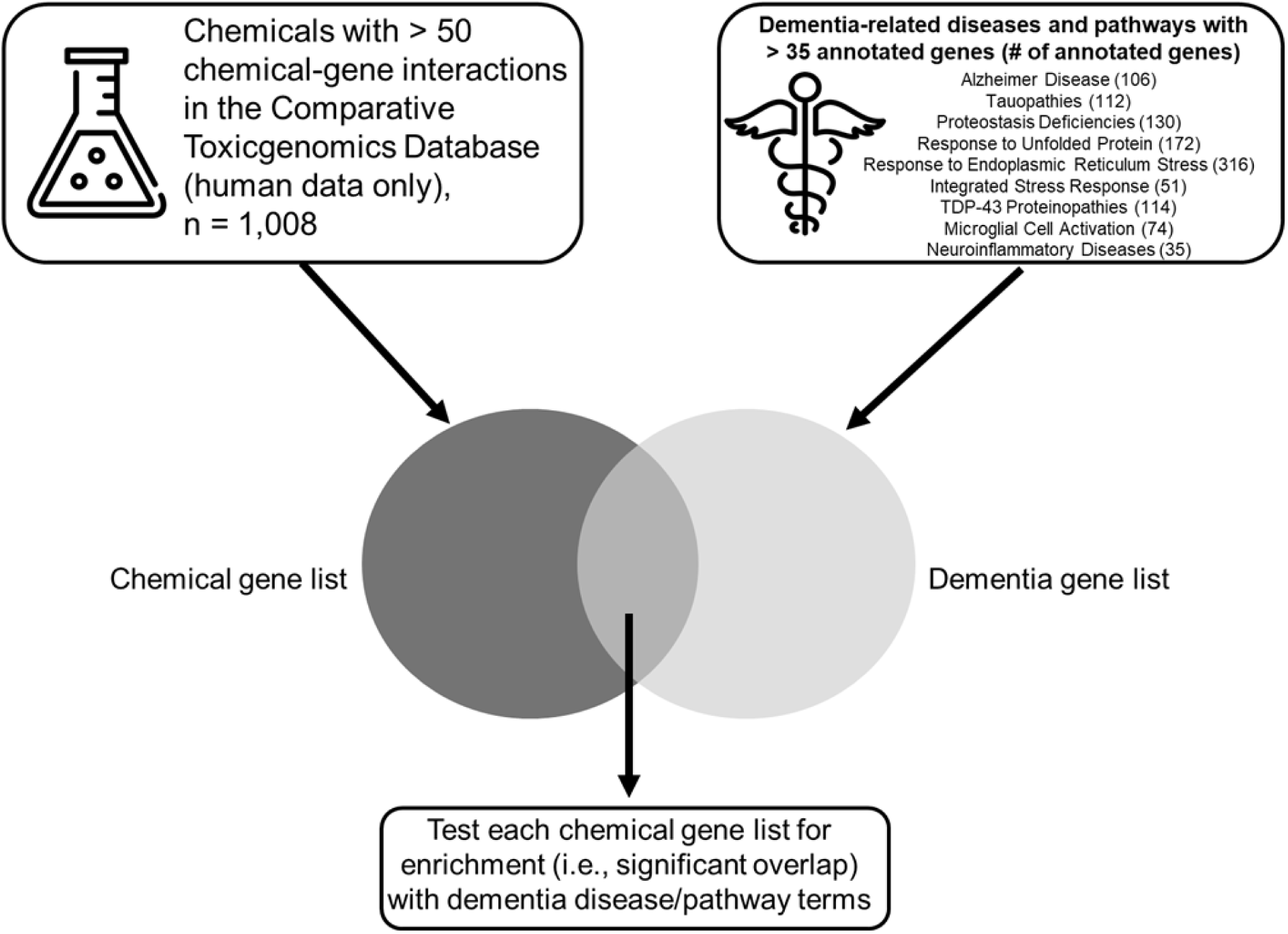
Summary of method and criteria for chemical and dementia disease gene lists used for enrichment testing. Chemicals with > 50 annotated chemical-gene interactions in the Comparative Toxicogenomics Database (CTD) were selected for enrichment testing using nine dementia-related disease or pathways terms. Gene lists for all dementia terms were also obtained from the CTD. Enrichment testing was conducted to determine which chemicals had a significantly high number of overlapping genes with each of the dementia gene lists (i.e., more overlapping genes between the chemical and dementia gene lists than would be expected by chance). Significant enrichment identified using FDR < 0.05 (Fisher’s exact test).

## 2. Methods

### 2.1 Background human gene list

Datasets were sourced from the CTD.^19^ We downloaded all genes (unique gene symbols) associated with the species *Homo sapiens* (human) from the CTD (date of download August 6, 2025). These 46,478 genes were used as the background gene list for enrichment testing.

### 2.2 Chemical gene lists

We used an agnostic approach for chemical selection, due to the large number of chemicals that have not yet been assessed for an association with dementia and associated molecular/cellular pathways. For all chemicals in the CTD database (n=15,167), we downloaded the list of curated chemical-gene interactions (date of download August 6, 2025, **Supplemental Table 1**). It is important to note that while these chemicals are associated with the annotated genes, in the CTD the dose of the chemical and the direction of chemical effects on the gene (e.g. gene expression upregulation or downregulation) are not specified. We excluded chemicals with sparse gene annotations (less than 50 genes annotated), resulting in 1,008 chemicals included in the study. We used a flowchart to show chemicals considered for study inclusion (**Supplemental Figure 1**).

### 2.3 Chemical categorization

For the chemicals meeting our inclusion criteria, we developed a chemical categorization scheme that built on categories defined by Nguyen, et al. (based on chemical biomarker data reported in the National Health and Nutrition Examination Survey)^20^ and chemical categories annotated in the Medical Subject Headings (MeSH) terms provided by CTD. Each chemical was assigned to one of 25 categories based on chemical use (e.g., “Pesticides, Pesticide Synergists and Cholinesterase Inhibitors”), public health or nutritional relevance (e.g., “Vitamins and Dietary Components”) or molecular structure (e.g., “Heterocyclic Compounds”). The full list of chemical categories and the distribution of the number of genes annotated to chemicals in each category are shown in **Table 1**. Chemical categories were used to provide context and examine patterns in the results.

**Table 1.**
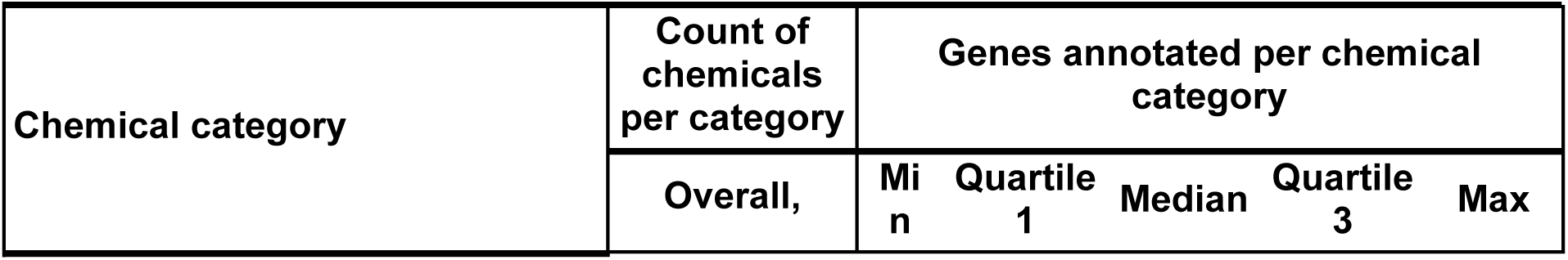
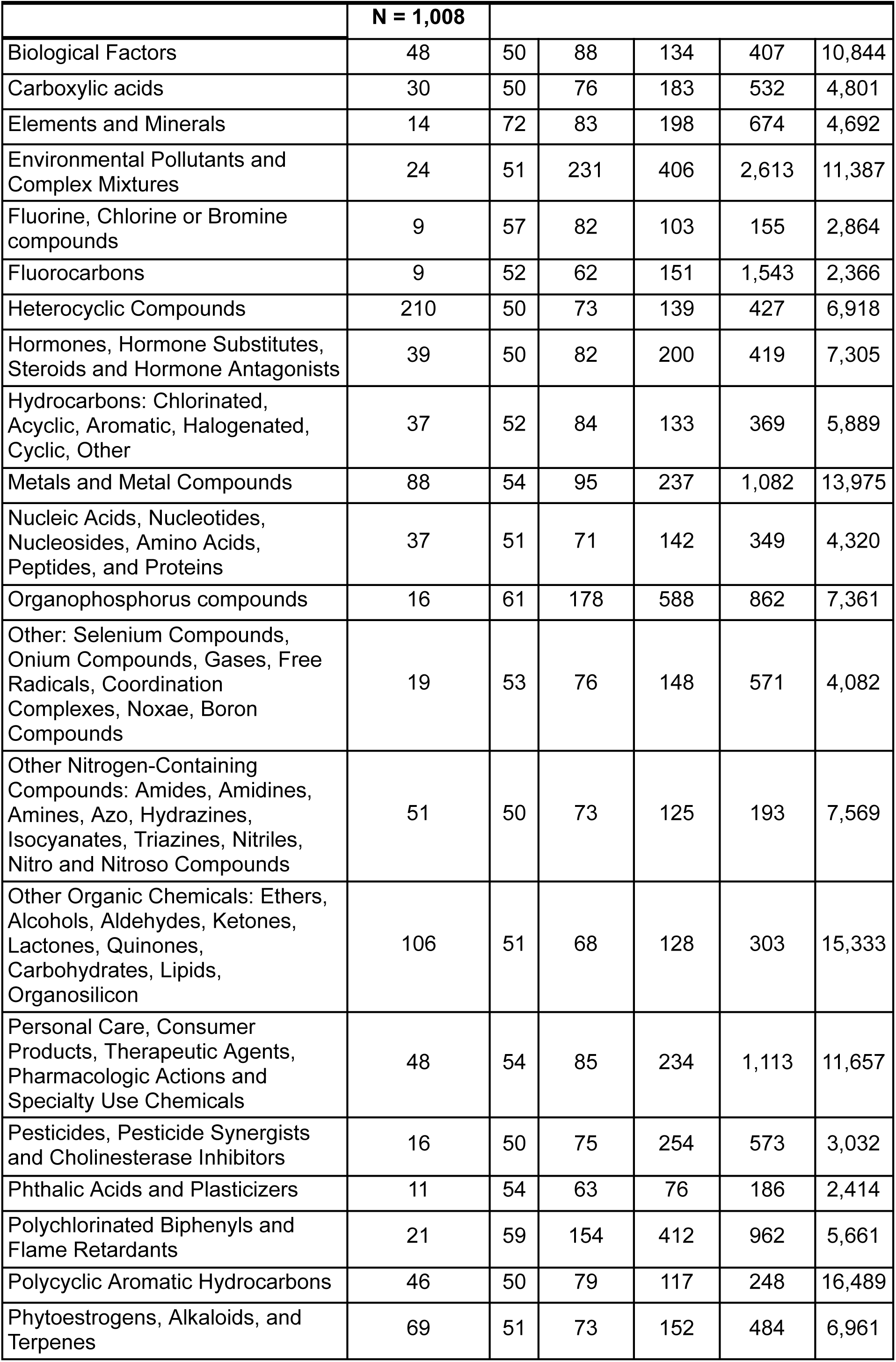
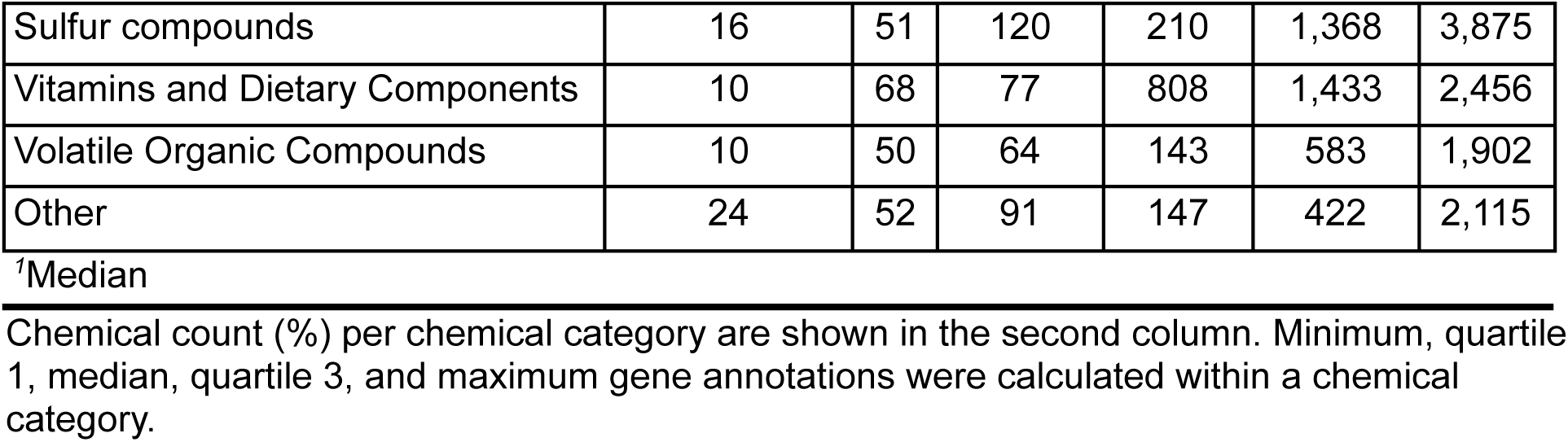
Descriptive statistics for the included chemical sample Number of chemicals per chemical category and distribution of genes per chemical category.

| Chemical category | Count of chemicals per category | Genes annotated per chemical category |  |  |  |  |
| --- | --- | --- | --- | --- | --- | --- |
|  | Overall, n | Min | Quartile 1 | Median | Quartile 3 | Max |

|  | **N = 1,008** |  |  |  |  |  |
| --- | --- | --- | --- | --- | --- | --- |
| Biological Factors | 48 | 50 | 88 | 134 | 407 | 10,844 |
| Carboxylic acids | 30 | 50 | 76 | 183 | 532 | 4,801 |
| Elements and Minerals | 14 | 72 | 83 | 198 | 674 | 4,692 |
| Environmental Pollutants and Complex Mixtures | 24 | 51 | 231 | 406 | 2,613 | 11,387 |
| Fluorine, Chlorine or Bromine compounds | 9 | 57 | 82 | 103 | 155 | 2,864 |
| Fluorocarbons | 9 | 52 | 62 | 151 | 1,543 | 2,366 |
| Heterocyclic Compounds | 210 | 50 | 73 | 139 | 427 | 6,918 |
| Hormones, Hormone Substitutes, Steroids and Hormone Antagonists | 39 | 50 | 82 | 200 | 419 | 7,305 |
| Hydrocarbons: Chlorinated, Acyclic, Aromatic, Halogenated, Cyclic, Other | 37 | 52 | 84 | 133 | 369 | 5,889 |
| Metals and Metal Compounds | 88 | 54 | 95 | 237 | 1,082 | 13,975 |
| Nucleic Acids, Nucleotides, Nucleosides, Amino Acids, Peptides, and Proteins | 37 | 51 | 71 | 142 | 349 | 4,320 |
| Organophosphorus compounds | 16 | 61 | 178 | 588 | 862 | 7,361 |
| Other: Selenium Compounds, Onium Compounds, Gases, Free Radicals, Coordination Complexes, Noxae, Boron Compounds | 19 | 53 | 76 | 148 | 571 | 4,082 |
| Other Nitrogen-Containing Compounds: Amides, Amidines, Amines, Azo, Hydrazines, Isocyanates, Triazines, Nitriles, Nitro and Nitroso Compounds | 51 | 50 | 73 | 125 | 193 | 7,569 |
| Other Organic Chemicals: Ethers, Alcohols, Aldehydes, Ketones, Lactones, Quinones, Carbohydrates, Lipids, Organosilicon | 106 | 51 | 68 | 128 | 303 | 15,333 |
| Personal Care, Consumer Products, Therapeutic Agents, Pharmacologic Actions and Specialty Use Chemicals | 48 | 54 | 85 | 234 | 1,113 | 11,657 |
| Pesticides, Pesticide Synergists and Cholinesterase Inhibitors | 16 | 50 | 75 | 254 | 573 | 3,032 |
| Phthalic Acids and Plasticizers | 11 | 54 | 63 | 76 | 186 | 2,414 |
| Polychlorinated Biphenyls and Flame Retardants | 21 | 59 | 154 | 412 | 962 | 5,661 |
| Polycyclic Aromatic Hydrocarbons | 46 | 50 | 79 | 117 | 248 | 16,489 |
| Phytoestrogens, Alkaloids, and Terpenes | 69 | 51 | 73 | 152 | 484 | 6,961 |
| Sulfur compounds | 16 | 51 | 120 | 210 | 1,368 | 3,875 |
| Vitamins and Dietary Components | 10 | 68 | 77 | 808 | 1,433 | 2,456 |
| Volatile Organic Compounds | 10 | 50 | 64 | 143 | 583 | 1,902 |
| Other | 24 | 52 | 91 | 147 | 422 | 2,115 |

<sup>1</sup>Median
Chemical count (%) per chemical category are shown in the second column. Minimum, quartile 1, median, quartile 3, and maximum gene annotations were calculated within a chemical category.

### 2.4 Dementia gene lists

Based on the known etiology and identified molecular/cellular hallmarks of dementia,^5,21–24^ we considered 12 disease terms or biological processes annotated in the CTD as relevant for evaluation: “Alzheimer Disease” (MESH: D000544), “Frontotemporal Dementia” (MeSH: D057180), “Tauopathies” (i.e., abnormal accumulation of tau proteins in the brain,^25^ a hallmark of AD; MeSH: D024801), “Lewy Body Disease” (MeSH: D020961), “Proteostasis Deficiences” (MeSH: D057165), “Lysosomal Storage Diseases, Nervous System” (MeSH: D020140), “TDP-43 Proteinopathies” (i.e., mislocalization of the Transactive response DNA binding protein of 43 kDa, characteristic of neurodegenerative diseases, including dementia,^26^ MeSH: D057177), “Neuroinflammatory Diseases” (MeSH: D000090862), “Response to Unfolded Protein” (GO:0006986), “Response to ER Stress” (GO:0034976), “Integrated Stress Response Signaling” (GO:0140467) and “Microglial Cell Activation” (GO:0001774). For each term, we downloaded the corresponding CTD gene list (date of download August, 6 2025, **Supplemental Table 2**). We excluded sparsely annotated terms and restricted our disease/pathways to those with a minimum gene set size of 35. We used a flowchart to show dementia terms considered for study inclusion (**Supplemental Figure 1**).

### 2.5 Testing chemical gene lists for enrichment with dementia and pathways

Analyses were performed using R statistical software (version 4.4.1).^27^ The code used to conduct these analyses and generate figures is publicly available (https://github.com/bakulskilab/CTD-dementia). We calculated univariate descriptive statistics for our chemical analytic sample including median, 25th percentile, 75th percentile, and interquartile range (IQR). Chemical count and percent were calculated by chemical category. Quartiles were calculated for gene annotations within a chemical category. We visualized the overlap of gene signatures in our dementia disease list pathways in an upset plot.

For our primary analyses, we generated pairwise 2 x 2 descriptive tables for chemical gene annotations by disease genes annotations. Based on these observed frequencies, we conducted enrichment tests between each chemical gene annotation and each dementia pathway gene annotation using Fisher’s exact tests. In situations where an observed 2 x 2 table returned a cell count of zero, we applied Laplace smoothing methodology, which adds one to each cell count.^28^ We accounted for multiple comparisons with the Benjamini-Hochberg method to calculate false discovery rate (FDR) adjusted p-values.^29^ We considered FDR < 0.05 to be nominally associated, and a threshold of FDR < x10^-6^ to be significantly associated. To determine the direction and magnitude of association, we calculated the proportional reporting ratio (PRR) with the 95% confidence interval (CI). A PRR value greater than one indicates enrichment (more overlapping genes than expected by chance), a PRR less than one indicates depletion (fewer overlapping genes than expected by chance), a PRR equal to one indicates neither. We visualized results across all dementia pathways using scatter plots of the magnitude of association and level of significance. Within each pathway, we ranked associations by smallest FDR. To consider results across pathways, we summed the ranked associations. In table format we presented the summed rank, number of genes annotated per chemical, rank within the disease list, PRR, and FDR. To visualize the results at the top ranked chemicals, we used heatmaps of PRR values by disease pathway and a forest plot of PRR values by disease pathway. To assess enrichment by chemical category, we viewed the number and percent of chemicals enriched by chemical category and disease list.

### 2.6 Assessing chemicals impacting key genes involved in neurodegenerative disease etiology

To gain insight into chemicals that impact key genes involved in the etiology of dementia, we quantified the number of curated chemical-gene interactions for five key genes involved in neurodegenerative disease etiology^6^: *APOE* (Apolipoprotein E), *APP* (amyloid-beta precursor protein), *MAPT* (microtubule associated protein tau), *PSEN1* (Presenilin-1) and *PSEN2* (Presenilin-2). All five of these genes and their protein products play important roles in the formation of the amyloid-beta plaques, one of the key pathological hallmarks of Alzheimer’s disease and genetic polymorphisms for these genes are associated with development of Alzheimer’s disease.^30–34^

## 3. Results

### 3.1 Chemical gene annotations

After filtering on the number of gene annotations, 1,008 chemicals from 25 chemical categories were included in the analysis (**Supplemental Figure 1A**). The chemical category containing the greatest number of chemicals was “heterocyclic compounds” (n = 210, 20.8%), including fluorescein-5-isocyanate, salinomycin and quinoline, and these chemicals were annotated to a median of 134 genes (IQR = 354) (**Supplemental Table 1**). The categories with the smallest number of chemicals were “fluorine, chlorine, or bromine compounds” (n = 9, 0.8%), including sodium fluoride, which were annotated to a median of 103 genes (IQR = 73), and “fluorocarbons” (n = 9, 0.8%), including perfluorohexanesulfonic acid, which were annotated to a median of 515 genes (IQR = 1,481).

### 3.2 Disease gene annotations

After applying filters for minimum number of gene annotations, nine disease lists were included in the analysis (**Supplemental Figure 1B**). “Response to ER Stress” had the largest gene set size (n = 348), while “Neuroinflammatory Diseases” had the smallest (n = 35). Across all disease pathways, the majority of genes were unique to a pathway (**Supplemental Figure 2**). The “Response to ER Stress” pathway had the most unique genes (i.e., genes not annotated to any of the other dementia disease terms, n = 199, 57.2%), while “Tauopathies” had the fewest (n = 3, 2.7%). The largest intersection (n = 22 genes) between gene lists was observed between “Response to ER Stress” (7.0%), “Response to Unfolded Protein” (12.8%), and “Integrated Stress Response” (19.6%).

### 3.3 Overall chemical gene associations with dementia pathways

In general, we observed extensive associations between the chemical gene lists and genes in the dementia pathways (**Figure 2**, **Supplemental Table 3**). Those associations were more likely to be enriched (PRR>1, in red in Figure 2) with more overlap than we would expect by chance, rather than depleted (PRR<1, in blue in Figure 2) with less overlap than we would expect by chance. When looking at pairwise associations between each chemical (n = 1,008) and disease list tested (n = 9), the greatest numbers of chemicals were significantly associated (FDR < 1×10^-6^) with the terms “Alzheimer’s disease” (n = 509) and “Tauopathies” (n = 511), while “Neuroinflammatory Diseases” had the least amount of associations in our chemical set (n = 75). If considering a nominal association (FDR < 0.05), we observed that between 90.3% (“Tauopathies”, n = 910 chemicals) and 66.1% (“Integrated Stress Response”, n = 666 chemicals) of the chemicals tested (n = 1,008), were associated with a dementia pathway.

**Figure 2.**
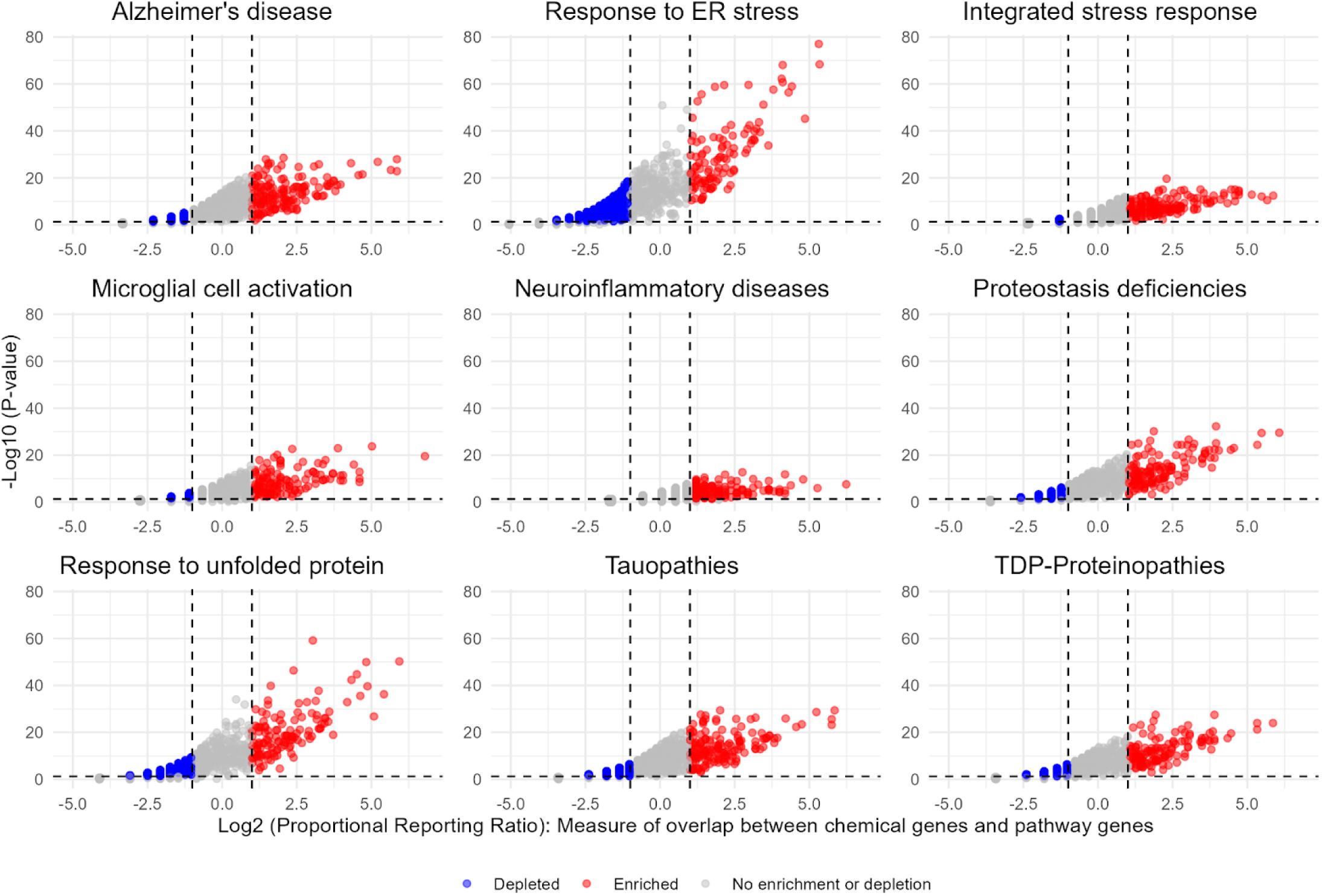
Scatter plots showing enrichment or depletion of chemical gene annotations for nine dementia pathways. Points in red represent enrichment, blue represents depletion, and gray represents no enrichment or depletion. The x-axis is log2 proportional reporting ratio (PRR). The y-axis is -log10 FDR-corrected p-value for Fisher’s Exact test for chemical gene annotation enrichment with each dementia pathway.

**Figure 3.**
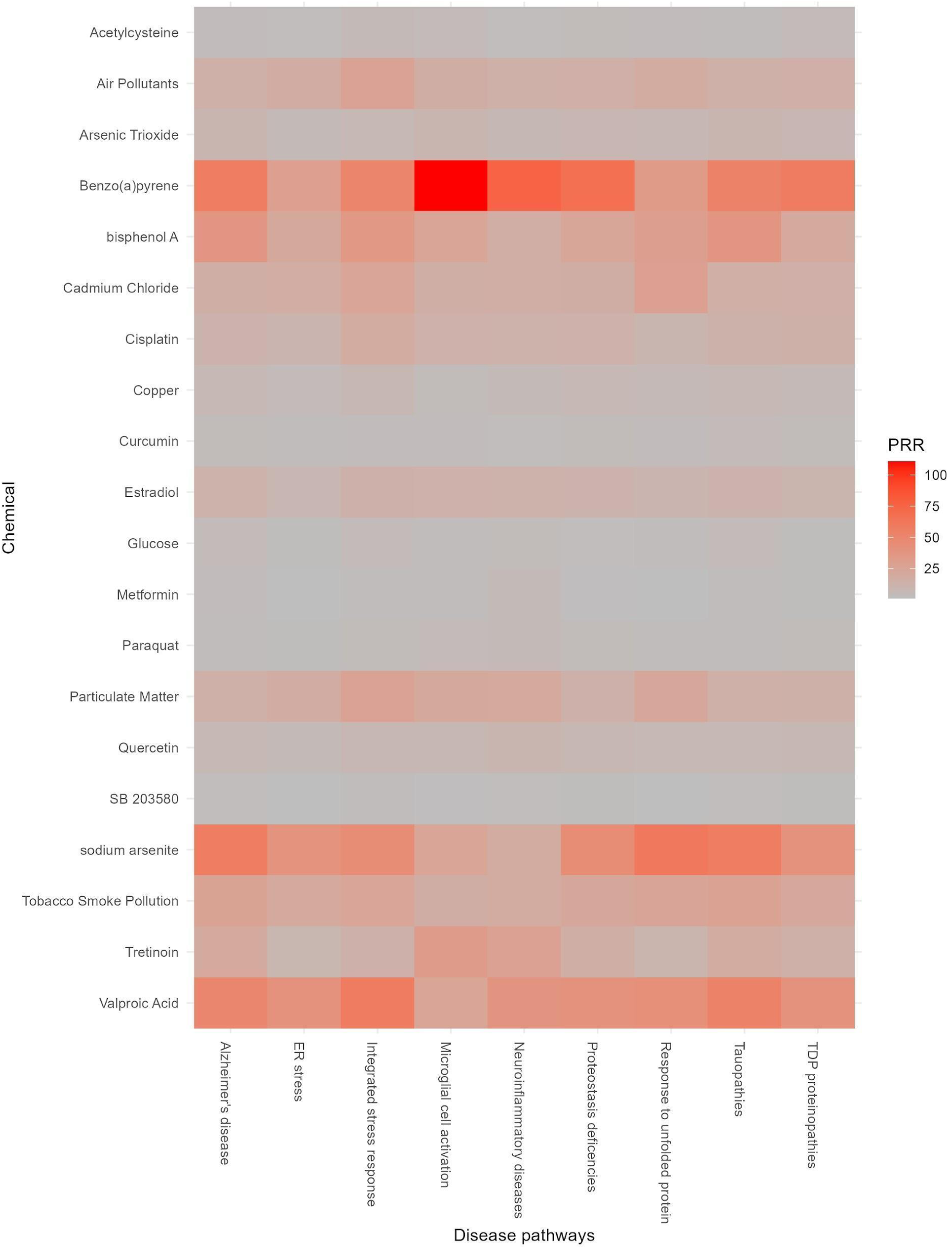
Heatmap of proportional reporting ratio (PRR) gene enrichment of top hits for nine dementia disease/pathways. Rows indicate chemicals tested and columns indicate the disease/pathway tested. The color of the cell indicates the PRR between the given chemical and disease/pathway, with red indicating a PRR closer to 100 and gray indicating a PRR closer to 1.

Out of the 1,008 chemicals tested, 396 (39.3%) were nominally enriched (FDR<0.05) across all nine dementia terms. Fifteen chemicals (1.5%) were enriched at FDR < 10^-6^ across all nine dementia terms (**Supplemental Table 4**). The top chemicals by summed rank across all nine disease lists included structurally diverse xenobiotic environmental toxicants such as benzo(a)pyrene, bisphenol A, cadmium, and arsenic, as well as endogenous compounds like glucose and estradiol (**Table 2, Supplemental Figure 3**). The greatest percentage (21.6%) of chemicals enriched across all nine dementia pathways was found in the chemical category of heterocyclic compounds (**Supplemental Table 5**).

**Table 2.**
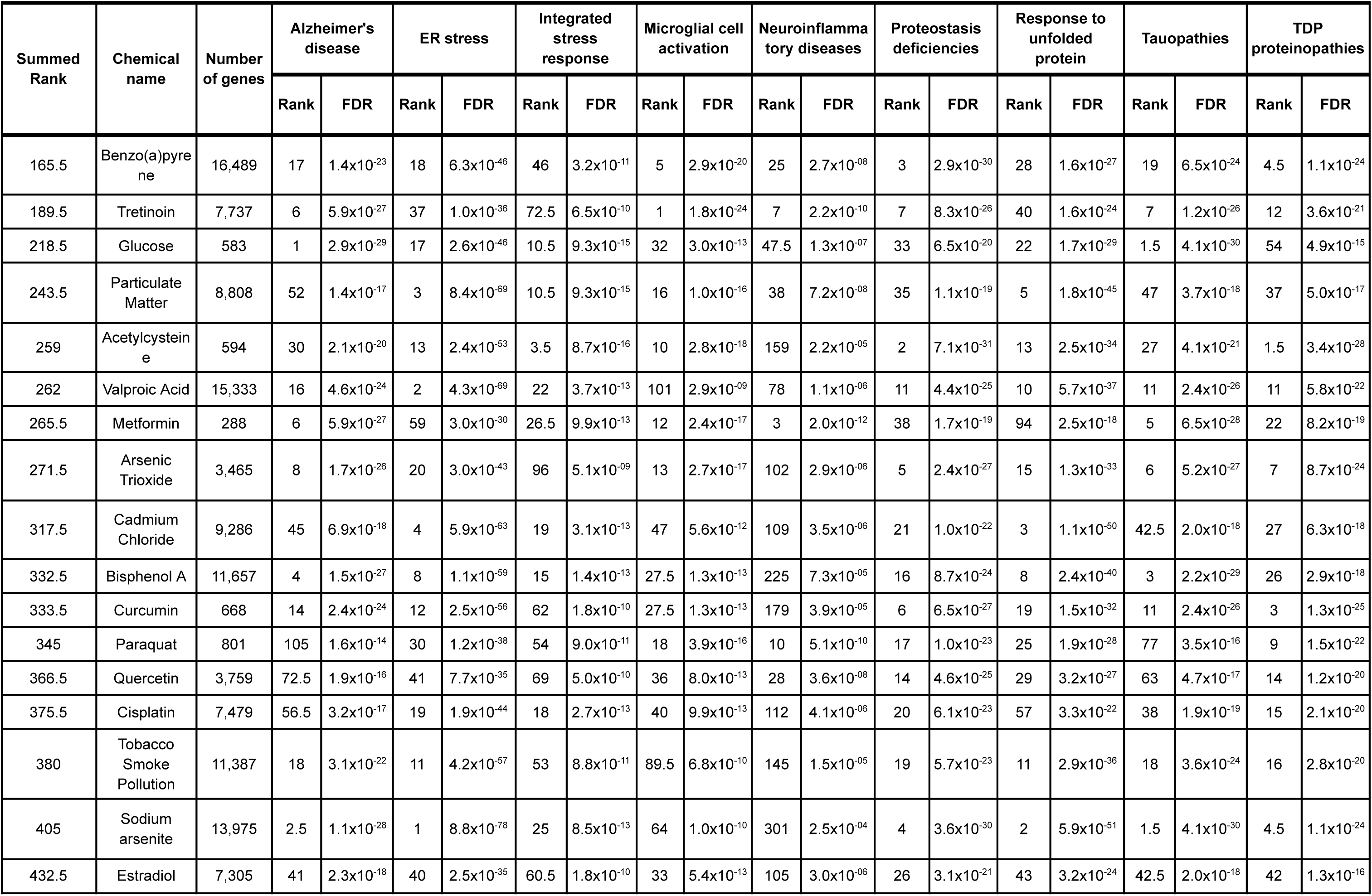
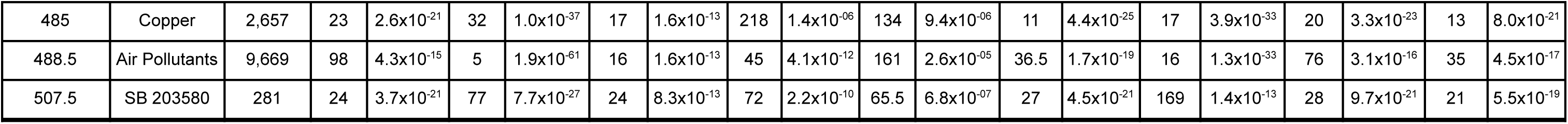
Table of the top 20 chemical gene annotations enriched for dementia pathways based on the summed rank of the minimum significance value.

| Summed Rank | Chemical name | Number of genes | Alzheimer's disease |  | ER stress |  | Integrated stress response |  | Microglial cell activation |  | Neuroinflammatory diseases |  | Proteostasis deficiencies |  | Response to unfolded protein |  | Tauopathies |  | TDP proteinopathies |  |
| --- | --- | --- | --- | --- | --- | --- | --- | --- | --- | --- | --- | --- | --- | --- | --- | --- | --- | --- | --- | --- |
|  |  |  | Rank | FDR | Rank | FDR | Rank | FDR | Rank | FDR | Rank | FDR | Rank | FDR | Rank | FDR | Rank | FDR | Rank | FDR |
| 165.5 | Benzo(a)pyrene | 16,489 | 17 | $1.4 \times 10^{-23}$ | 18 | $6.3 \times 10^{-46}$ | 46 | $3.2 \times 10^{-11}$ | 5 | $2.9 \times 10^{-20}$ | 25 | $2.7 \times 10^{-08}$ | 3 | $2.9 \times 10^{-30}$ | 28 | $1.6 \times 10^{-27}$ | 19 | $6.5 \times 10^{-24}$ | 4.5 | $1.1 \times 10^{-24}$ |
| 189.5 | Tretinoin | 7,737 | 6 | $5.9 \times 10^{-27}$ | 37 | $1.0 \times 10^{-36}$ | 72.5 | $6.5 \times 10^{-10}$ | 1 | $1.8 \times 10^{-24}$ | 7 | $2.2 \times 10^{-10}$ | 7 | $8.3 \times 10^{-26}$ | 40 | $1.6 \times 10^{-24}$ | 7 | $1.2 \times 10^{-26}$ | 12 | $3.6 \times 10^{-21}$ |
| 218.5 | Glucose | 583 | 1 | $2.9 \times 10^{-29}$ | 17 | $2.6 \times 10^{-46}$ | 10.5 | $9.3 \times 10^{-15}$ | 32 | $3.0 \times 10^{-13}$ | 47.5 | $1.3 \times 10^{-07}$ | 33 | $6.5 \times 10^{-20}$ | 22 | $1.7 \times 10^{-29}$ | 1.5 | $4.1 \times 10^{-30}$ | 54 | $4.9 \times 10^{-15}$ |
| 243.5 | Particulate Matter | 8,808 | 52 | $1.4 \times 10^{-17}$ | 3 | $8.4 \times 10^{-69}$ | 10.5 | $9.3 \times 10^{-15}$ | 16 | $1.0 \times 10^{-16}$ | 38 | $7.2 \times 10^{-08}$ | 35 | $1.1 \times 10^{-19}$ | 5 | $1.8 \times 10^{-45}$ | 47 | $3.7 \times 10^{-18}$ | 37 | $5.0 \times 10^{-17}$ |
| 259 | Acetylcysteine | 594 | 30 | $2.1 \times 10^{-20}$ | 13 | $2.4 \times 10^{-53}$ | 3.5 | $8.7 \times 10^{-16}$ | 10 | $2.8 \times 10^{-18}$ | 159 | $2.2 \times 10^{-05}$ | 2 | $7.1 \times 10^{-31}$ | 13 | $2.5 \times 10^{-34}$ | 27 | $4.1 \times 10^{-21}$ | 1.5 | $3.4 \times 10^{-28}$ |
| 262 | Valproic Acid | 15,333 | 16 | $4.6 \times 10^{-24}$ | 2 | $4.3 \times 10^{-69}$ | 22 | $3.7 \times 10^{-13}$ | 101 | $2.9 \times 10^{-09}$ | 78 | $1.1 \times 10^{-06}$ | 11 | $4.4 \times 10^{-25}$ | 10 | $5.7 \times 10^{-37}$ | 11 | $2.4 \times 10^{-26}$ | 11 | $5.8 \times 10^{-22}$ |
| 265.5 | Metformin | 288 | 6 | $5.9 \times 10^{-27}$ | 59 | $3.0 \times 10^{-30}$ | 26.5 | $9.9 \times 10^{-13}$ | 12 | $2.4 \times 10^{-17}$ | 3 | $2.0 \times 10^{-12}$ | 38 | $1.7 \times 10^{-19}$ | 94 | $2.5 \times 10^{-18}$ | 5 | $6.5 \times 10^{-28}$ | 22 | $8.2 \times 10^{-19}$ |
| 271.5 | Arsenic Trioxide | 3,465 | 8 | $1.7 \times 10^{-26}$ | 20 | $3.0 \times 10^{-43}$ | 96 | $5.1 \times 10^{-09}$ | 13 | $2.7 \times 10^{-17}$ | 102 | $2.9 \times 10^{-06}$ | 5 | $2.4 \times 10^{-27}$ | 15 | $1.3 \times 10^{-33}$ | 6 | $5.2 \times 10^{-27}$ | 7 | $8.7 \times 10^{-24}$ |
| 317.5 | Cadmium Chloride | 9,286 | 45 | $6.9 \times 10^{-18}$ | 4 | $5.9 \times 10^{-63}$ | 19 | $3.1 \times 10^{-13}$ | 47 | $5.6 \times 10^{-12}$ | 109 | $3.5 \times 10^{-06}$ | 21 | $1.0 \times 10^{-22}$ | 3 | $1.1 \times 10^{-50}$ | 42.5 | $2.0 \times 10^{-18}$ | 27 | $6.3 \times 10^{-18}$ |
| 332.5 | Bisphenol A | 11,657 | 4 | $1.5 \times 10^{-27}$ | 8 | $1.1 \times 10^{-59}$ | 15 | $1.4 \times 10^{-13}$ | 27.5 | $1.3 \times 10^{-13}$ | 225 | $7.3 \times 10^{-05}$ | 16 | $8.7 \times 10^{-24}$ | 8 | $2.4 \times 10^{-40}$ | 3 | $2.2 \times 10^{-29}$ | 26 | $2.9 \times 10^{-18}$ |
| 333.5 | Curcumin | 668 | 14 | $2.4 \times 10^{-24}$ | 12 | $2.5 \times 10^{-56}$ | 62 | $1.8 \times 10^{-10}$ | 27.5 | $1.3 \times 10^{-13}$ | 179 | $3.9 \times 10^{-05}$ | 6 | $6.5 \times 10^{-27}$ | 19 | $1.5 \times 10^{-32}$ | 11 | $2.4 \times 10^{-26}$ | 3 | $1.3 \times 10^{-25}$ |
| 345 | Paraquat | 801 | 105 | $1.6 \times 10^{-14}$ | 30 | $1.2 \times 10^{-38}$ | 54 | $9.0 \times 10^{-11}$ | 18 | $3.9 \times 10^{-16}$ | 10 | $5.1 \times 10^{-10}$ | 17 | $1.0 \times 10^{-23}$ | 25 | $1.9 \times 10^{-28}$ | 77 | $3.5 \times 10^{-16}$ | 9 | $1.5 \times 10^{-22}$ |
| 366.5 | Quercetin | 3,759 | 72.5 | $1.9 \times 10^{-16}$ | 41 | $7.7 \times 10^{-35}$ | 69 | $5.0 \times 10^{-10}$ | 36 | $8.0 \times 10^{-13}$ | 28 | $3.6 \times 10^{-08}$ | 14 | $4.6 \times 10^{-25}$ | 29 | $3.2 \times 10^{-27}$ | 63 | $4.7 \times 10^{-17}$ | 14 | $1.2 \times 10^{-20}$ |
| 375.5 | Cisplatin | 7,479 | 56.5 | $3.2 \times 10^{-17}$ | 19 | $1.9 \times 10^{-44}$ | 18 | $2.7 \times 10^{-13}$ | 40 | $9.9 \times 10^{-13}$ | 112 | $4.1 \times 10^{-06}$ | 20 | $6.1 \times 10^{-23}$ | 57 | $3.3 \times 10^{-22}$ | 38 | $1.9 \times 10^{-19}$ | 15 | $2.1 \times 10^{-20}$ |
| 380 | Tobacco Smoke Pollution | 11,387 | 18 | $3.1 \times 10^{-22}$ | 11 | $4.2 \times 10^{-57}$ | 53 | $8.8 \times 10^{-11}$ | 89.5 | $6.8 \times 10^{-10}$ | 145 | $1.5 \times 10^{-05}$ | 19 | $5.7 \times 10^{-23}$ | 11 | $2.9 \times 10^{-36}$ | 18 | $3.6 \times 10^{-24}$ | 16 | $2.8 \times 10^{-20}$ |
| 405 | Sodium arsenite | 13,975 | 2.5 | $1.1 \times 10^{-28}$ | 1 | $8.8 \times 10^{-78}$ | 25 | $8.5 \times 10^{-13}$ | 64 | $1.0 \times 10^{-10}$ | 301 | $2.5 \times 10^{-04}$ | 4 | $3.6 \times 10^{-30}$ | 2 | $5.9 \times 10^{-51}$ | 1.5 | $4.1 \times 10^{-30}$ | 4.5 | $1.1 \times 10^{-24}$ |
| 432.5 | Estradiol | 7,305 | 41 | $2.3 \times 10^{-18}$ | 40 | $2.5 \times 10^{-35}$ | 60.5 | $1.8 \times 10^{-10}$ | 33 | $5.4 \times 10^{-13}$ | 105 | $3.0 \times 10^{-06}$ | 26 | $3.1 \times 10^{-21}$ | 43 | $3.2 \times 10^{-24}$ | 42.5 | $2.0 \times 10^{-18}$ | 42 | $1.3 \times 10^{-16}$ |
| 485 | Copper | 2,657 | 23 | $2.6 \times 10^{-21}$ | 32 | $1.0 \times 10^{-37}$ | 17 | $1.6 \times 10^{-13}$ | 218 | $1.4 \times 10^{-06}$ | 134 | $9.4 \times 10^{-06}$ | 11 | $4.4 \times 10^{-25}$ | 17 | $3.9 \times 10^{-33}$ | 20 | $3.3 \times 10^{-23}$ | 13 | $8.0 \times 10^{-21}$ |
| 488.5 | Air Pollutants | 9,669 | 98 | $4.3 \times 10^{-15}$ | 5 | $1.9 \times 10^{-61}$ | 16 | $1.6 \times 10^{-13}$ | 45 | $4.1 \times 10^{-12}$ | 161 | $2.6 \times 10^{-05}$ | 36.5 | $1.7 \times 10^{-19}$ | 16 | $1.3 \times 10^{-33}$ | 76 | $3.1 \times 10^{-16}$ | 35 | $4.5 \times 10^{-17}$ |
| 507.5 | SB 203580 | 281 | 24 | $3.7 \times 10^{-21}$ | 77 | $7.7 \times 10^{-27}$ | 24 | $8.3 \times 10^{-13}$ | 72 | $2.2 \times 10^{-10}$ | 65.5 | $6.8 \times 10^{-07}$ | 27 | $4.5 \times 10^{-21}$ | 169 | $1.4 \times 10^{-13}$ | 28 | $9.7 \times 10^{-21}$ | 21 | $5.5 \times 10^{-19}$ |

### 3.4 Chemical gene enrichment with individual dementia pathways

Of the chemicals associated with “Alzheimer’s Disease” genes, we observed 58.1% (295 chemicals) were enriched, meaning that they had more overlapping genes than would be expected by chance at FDR<10^-6^ (**Supplemental Table 5**). Among the enriched chemicals, 16.3% were chemicals from the heterocyclic compounds category and 12.2% were metals or metalloids. The chemicals most significantly enriched with “Alzheimer’s Disease” genes based on FDR were glucose (PRR = 4.2), sodium arsenite (PRR = 57.9), and chlorpyrifos (PRR = 2.8). We saw that 70% of vitamins and dietary components tested were associated with “Alzheimer’s Disease”.

For the “Response to ER Stress” pathway, 33.8% of associations were enriched (200 chemicals). Of the enriched chemicals, 17.5% were heterocyclic compounds and 7.5% were personal care and consumer products. The most significantly enriched chemicals were sodium arsenite (PRR=39.9), valproic acid (PRR=40.6), and particulate matter (PRR=17.2).

We found 193 chemicals associated with the “Integrated Stress Response” pathway and all were enriched (none were depleted). The most significantly enriched chemicals were benzyloxycarbonylleucyl-leucyl-leucine aldehyde (PRR=4.9), triphenyl phosphate (PRR=24.1.0), and abrine (PRR=22.0). We saw 42.1% of all selenium compounds tested were enriched for the “Integrated Stress Response” pathway.

For the “Microglial Cell Activation” pathway, we observed 213 chemical associations and 80.8% of those associations were enriched. The most significantly enriched chemicals were tretinoin (PRR=32.4), nickel (PRR=14.7), and tetradecanoylphorbol acetate (PRR=5.1). We observed 42.9% of all elements and minerals tested and 33.3% of all complex mixtures tested were enriched for genes in the “Microglial Cell Activation” pathway.

We observed 75 chemicals associated with the “Neuroinflammatory Diseases” pathway, and similar to the “Integrated Stress Response” pathway, all were enriched. Of the enriched chemicals, 17% were heterocyclic compounds and 12% were biologic factors. The most significantly enriched chemicals were tetradecanoylphorbol acetate (PRR=6.8), nickel (PRR=18.2), and metformin (PRR=4.5).

For the “Proteostasis Deficiencies” pathway, we observed 414 chemical associations and 55.8% of associations were enriched (231 chemicals). The most significantly enriched chemicals were resveratrol (PRR=15.6), acetylcysteine (PRR=3.7), and benzo(a)pyrene (PRR=67.3). We saw 60% of all vitamins and dietary components tested, 45.8% of complex mixtures tested, and 37.5% of pesticides tested were enriched for genes in the “Proteostasis Deficiencies” pathway.

Of the 404 chemical associations with the “Response to Unfolded Proteins” pathway, 56.9% were enriched. The most significantly enriched chemicals were tunicamycin (PRR=8.2), sodium arsenite (PRR=61.2), and cadmium chloride (PRR=28.4). Similar to the results from the “Proteostasis Deficiencies” pathway, for “Response to Unfolded Proteins”, we observed that 60% of all vitamins and dietary components tested, 45.8% of complex mixtures tested, and 36.8% of selenium compounds were enriched for genes in the “Response to Unfolded Proteins” pathway.

For the “Tauopathies” pathway, we observed 511 associations and 59.7% of those chemicals were enriched for genes in the “Tauopathies” pathway. The most significantly enriched chemicals were sodium arsenite (PRR=57.5), glucose (PRR=4.0), and bisphenol A (PRR=37.7). We observed 70% of all vitamins and dietary components tested, 42.9% of complex mixtures, and 42.1% of selenium compounds tested were enriched for genes in the “Tauopathies” pathway.

We observed 476 chemicals associated with the “TDP-proteinopathies” pathway, and 67.4% of these associations were enriched. The most significantly enriched chemicals were acetylcysteine (PRR=3.8), resveratrol (PRR=14.9), and curcumin (PRR=3.6). We saw 60% of all vitamins and dietary components tested, 42.9% of elements and minerals, and 41.7% of complex mixtures tested were enriched for genes in the “TDP-proteinopathies” pathway.

### 3.5 Key Alzheimer’s disease genes impacted by chemicals

We examined five key genes involved in neurodegenerative disease etiology (*APOE*, *APP*, *MAPT*, *PSEN1*, and *PSEN2*) for appearance in chemical gene annotations (**Table 3**). *APP* was observed in the gene annotations for 163 chemicals (16%), including glucose and lead. The presenilin genes were least represented in chemical gene annotations. *PSEN1* was observed in 52 chemical gene lists (5% of chemicals tested) and *PSEN2* was observed in 42 gene lists (4% of chemicals tested). The full list of chemical-gene annotations for the five key Alzheimer’s disease genes is shown in **Supplemental Table 6**.

**Table 3.** Table showing number (%) chemicals in the analytic sample (N=1,008) that impact specific dementia hallmark genes.

| Alzheimer's disease Dementia hallmark genes | Number of chemicals | Example chemicals relevant to human exposome |
| --- | --- | --- |
| *APOE* | 109 (11%) | Aflatoxin B1, Arsenic, Benzo(a)pyrene, Lead, Perfluorooctane Sulfonic Acid |
| *APP* | 163 (16%) | Benzene, Bisphenol A, Glyphosate, Perfluorobutanesulfonic Acid, Perfluorooctanoic Acid |
| *PSEN1* | 52 (5%) | Atrazine, Benzo(a)pyrene, Lead, Particulate Matter, Vehicle Emissions |
| *PSEN2* | 42 (4%) | Arsenic, Bisphenol A, Cannabidiol, Glyphosate, Valproic Acid |
| *MAPT* | 113 (11%) | Aflatoxin B1, Arsenic, Methamphetamine, Particulate Matter, Tobacco Smoke Pollution |

## 4. Discussion

Dementia is an impactful disorder and environmental exposures are modifiable risk factors.^35–40^ Given the large number of chemicals in the modern human exposome, as well as gaps in our understanding of cellular and molecular mechanisms of toxicity, prioritizing chemicals for public health interventions is essential. Using the CTD, we screened a large and diverse set of chemicals for gene enrichment with dementia, observing 742 chemicals associated with at least one dementia pathway and 15 associated with all nine dementia pathways (2-(2-amino-3-methoxyphenyl)-4H-1-benzopyran-4-one, benzo(a)pyrene, capsaicin, ethanol, glucose, glutathione, lipopolysaccharides, metformin, paraquat, particulate matter, quercetin, rosiglitazone, SB 203580, tetradecanoylphorbol acetate, and tretinoin). This study provides important insights into chemicals warranting further dementia mechanistic evaluation and contributes understanding of how the environment may drive dementia risk.

After assessing 1,008 chemicals in 25 categories, we demonstrated diverse chemical structures and uses are linked to dementia-related molecular pathways. These findings build on recent chemical exposure-wide association studies of cognition among older adults.^41,42^ We found chemicals like cadmium, lead, and ozone were enriched for “Alzheimer’s Disease” genes, consistent with human epidemiology studies linking these exposures to dementia risk.^14,43^ Chemicals enriched for “Alzheimer’s Disease” are used in a variety of widespread industrial or agricultural processes, such as plasticizers (diethylhexyl phthalate) or pesticides (glyphosate), leading to possible exposures for manufacturing^44^ and agriculture^45^ workers, as well as consumers of products^46^ and residents near farms^47^ or manufacturing facilities.^48^ Workers in these industries are more likely to be from disadvantaged groups with less access to education, a known risk factor for dementia, and occupational exposures may be a hidden mechanism contributing to dementia among vulnerable populations.^49^ An individual’s total environmental exposure occurs via multiple pathways, including food and drink consumption, personal care product use, residential location, the workplace, and through recreational activities. Overall, we observed chemicals from all 25 categories were enriched for one or more dementia pathways, highlighting the diversity of molecular structures, chemical uses and potential exposure pathways for chemicals influencing dementia risk. Of the chemical categories assessed, “environmental pollutants and complex mixtures”, including tobacco smoke, particulate matter, and coal ash, had the highest percentage of chemicals enriched for dementia pathway genes, consistent with studies showing exposures to these pollutants increases risk for cognitive decline and dementia.^50–53^

Our study provides insight into pathways linking environmental exposures and dementia, including disruptions in protein homeostasis, activation of cell stress pathways, and activation of neuroinflammation. Protein homeostasis encompasses the controlled synthesis, folding, post-translational modification, and degradation of proteins, all of which are necessary for proper brain and neurological functions.^54^ Proteinopathy involves protein accumulation in the brain due to dysfunctions in protein homeostasis.^55^ We identified chemicals enriched with genes involved in proteinopathies consistent with dementia (“Tauopathies”, “TDP-43 Proteinopathies” and “Proteostasis Deficiences”). Toxicants disrupt protein homeostasis by altering cellular redox status, leading to free radical stress and interfering with protein folding,^56^ or by binding protein functional groups.^57^ These effects have been demonstrated with metals and pesticides,^56^ and our findings suggest a broader range of chemicals including polycyclic aromatic hydrocarbons like benzo(a)pyrene and pyrazolanthrone, fluorinated compounds like perfluorooctanoic acid, consumer product chemicals like bisphenol A, and endogenous chemicals like glucose (as in cases of poor glycemic control^58^) are linked to disrupted protein homeostasis.

We showed diverse chemicals were enriched with cellular stress response genes. In response to stressful events, these pathways affect protein homeostasis or inflammation (“Integrated Stress Response”, “Response to ER Stress”, and “Response to Unfolded Protein”). The Integrated Stress Response is an adaptive cellular pathway that restores homeostasis, including amino acid deprivation, viral infections, and reactive oxygen species.^59^ While cytoprotective when activated in the short term, prolonged or high intensity stress can cause the Integrated Stress Response pathway to promote cell death via apoptosis.^59,60^ A main activation signaling event is phosphorylation of the alpha subunit of the eukaryotic initiator factor 2 (eIF2α).^59^ Prolonged eIF2α phosphorylation in mice causes cognitive deficits.^6^ Our findings suggest environmental toxicants influence the Integrated Stress Response pathway, contributing to dementia. People may be chronically exposed to air pollutants or drinking water contaminants. If chemicals additively or synergistically activate the Integrated Stress Response pathway in a prolonged manner under chronic exposure conditions, these exposures could impact neurodegenerative diseases. Future studies should seek to understand the net effects on the Integrated Stress Response pathway in neurons in the context of dementia, particularly for the chemicals identified.

Prolonged activation of stress pathways and phosphorylated eIF2α in particular, can lead to neuroinflammation.^6^ We identified chemicals enriched for pathways related to neuroinflammation (“Neuroinflammation”, “Microglial Activation”). Neuroinflammation can lead to the death of neurons and damage to brain tissue.^61^ Alzheimer’s disease cases have increased inflammation in brain regions including the inferior and middle temporal gyri, left amygdala, and inferior parietal lobes.^61^ Our study demonstrated chemical classes including “environmental pollutants and complex mixtures”, “personal care”, “consumer products”, “therapeutic agents”, “pharmacologic actions and specialty use chemicals”, and “selenium compounds” impact neuroinflammation and microglial activation genes.

Our top hit chemicals by summed rank included xenobiotic environmental pollutants such as benzo(a)pyrene, bisphenol A, and arsenic compounds (e.g., sodium arsenite), consistent with epidemiology and animal studies linking these chemicals to neurodegenerative effects.^62–64^ Interestingly, top hit chemicals also included endogenous compounds like glucose and estradiol (the main estrogen hormone), and we emphasize we are unable to determine direction of gene activity in the CTD. Poor glycemic control^58^ and estrogen therapy^65^ have both been associated with dementia in human studies. However, protective effects have also been observed.^66^ We found an array of chemicals impact key neurodegenerative hallmark genes (*APOE*, *APP*, *PSEN1*, *PSEN2,* and *MAPT)*. Previous studies showed bisphenol A,^67^ lead,^68^ benzo(a)pyrene,^62^ and ochratoxin A^69^ alter expression of these genes, which we confirmed. A relatively low percentage of the chemicals assessed (4-16%) impacted each of these hallmark genes, suggesting many chemicals contribute to dementia risks through alternate gene interactions, requiring additional research.

We identified several dietary factors enriched with dementia pathways (ascorbic acid (vitamin C), vitamin E, selenium). We note our study was limited to identifying statistical associations between chemicals and dementia-relevant genes and the direction of chemical effects on genes was not specified.^19^ Thus, enrichment with dementia pathways for these dietary factors could indicate these compounds interacting with dementia pathways in a protective manner, and additional work is needed to determine the direction of these and all associations identified in the present manuscript. Maintaining adequate levels of, or supplementation with, dietary factors is protective against dementia^70–72^ and cognitive impairment, though findings from other meta-analyses or clinical trials have been mixed or negative, potentially due to impurities in supplements^73^ and highlighting the need for further study.^74^ Overall, our findings highlight the exposome’s complex impact on dementia risk. Future *in vitro* or *in vivo* mechanistic studies could draw on these findings to design chemical mixture studies to clarify the influences of toxicants, nutrients, and therapeutic compounds on dementia.

One important advantage of our methods is the ability to broadly screen a large number of chemicals for dementia-relevant associations to inform standard *in vitro* or *in vivo* toxicology models. We screened chemicals for multiple mechanisms of toxicity at once, including cellular stress responses, proteinopathies, and neuroinflammation. Limitations of this study should also be noted. Our screening efforts covered a considerable range of chemicals, however these represent only a portion of the chemicals (1,008 screened/15,167 present; 6.6%) in the CTD and in use globally. Significant gaps in our understanding of the broader chemical landscape and associations with dementia risk remain and require further research. Toxicology data from the CTD is not tissue or cell type-specific, meaning some of the assessed associations may not be relevant for neurons or glial cells. However, cell stress and inflammation are observed across a wide array of cell types, and chemicals can influence neurons indirectly by inducing stress in another cell type. Doses of chemicals may vary across studies annotated in the CTD and dose-responses for chemical-gene associations are not discernable. Finally, we tested chemicals for interactions with genes only. Direction and magnitude of gene expression changes was not assessed. To address these limitations, future studies should use *in vitro* or *in vivo* models to confirm associations between chemicals and the dementia pathways, prioritizing the genes identified in this study. For example, studies using neuronal or microglial cell models in high-throughput testing systems could screen plausible chemical mixtures for activation or inflammation markers or assess additive, synergistic, or counteracting effects on endoplasmic reticulum stress or integrated stress response pathways. Functional assays, including enzyme-linked immunosorbent assays targeting dementia pathology, electrophysiological assays capturing loss of synaptic function, and assays of mitochondrial dysfunction and oxidative stress (e.g., quantification of Aβ-Binding Alcohol Dehydrogenase or lipid peroxidation) would be useful expansions.

In conclusion, this hypothesis-generating screening study shows diverse chemicals overlap with molecular and cellular pathways consistent with the development of dementia. These chemicals may act through mechanisms including activation of cellular stress pathways and neuroinflammation. Given the significant challenges of understanding the exposome’s impact on the risk for dementia, these findings provide important information to inform future epidemiology and toxicology studies seeking to understand the link between environmental exposures and dementia.

## Sources of funding

This research was supported by the National Institute on Aging (R01 AG067592, R01 AG070897, P30 AG072931) and the National Institute of Environmental Health Sciences (P42 ES017198).

### Acknowledgements

We appreciate the staff of the Comparative Toxicogenomics Database.

## Data availability

All data used for this analysis are publicly available and can be downloaded from the Comparative Toxicogenomics Database webpage at https://ctdbase.org/.

