## Supplemental material for "Identification of chemicals targeting dementia genes and pathways in the Comparative Toxicogenomics Database"

**Supplementary Material**

This document contains three supplemental figures and six supplemental tables.

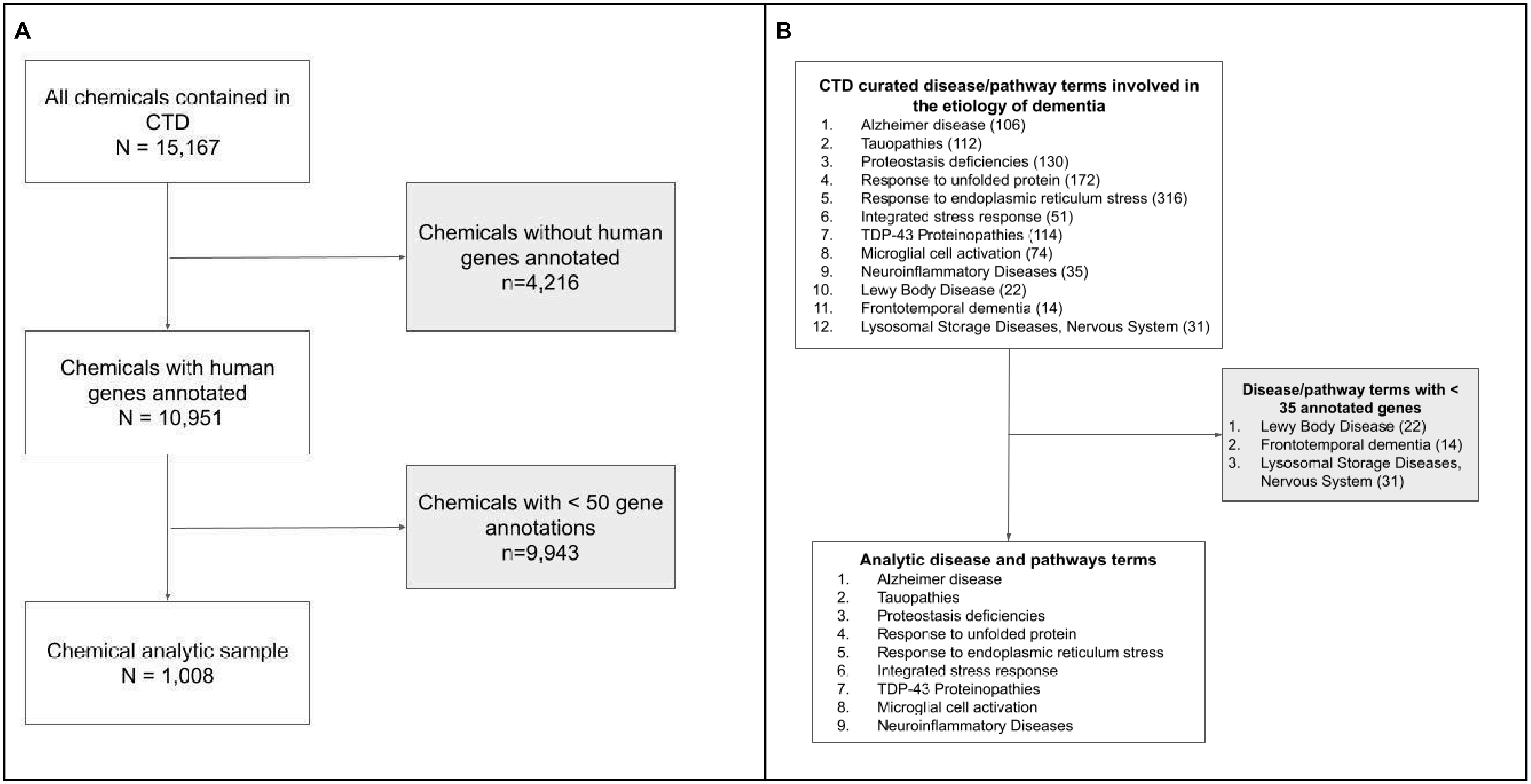


**Supplemental Figure 1.** Flowchart for study inclusion. Panel A represents the chemical selection. Panel B represents the selection for disease/pathway terms related to dementia etiology. For panel B, the number of genes annotated to each disease or pathway are shown in parentheses.

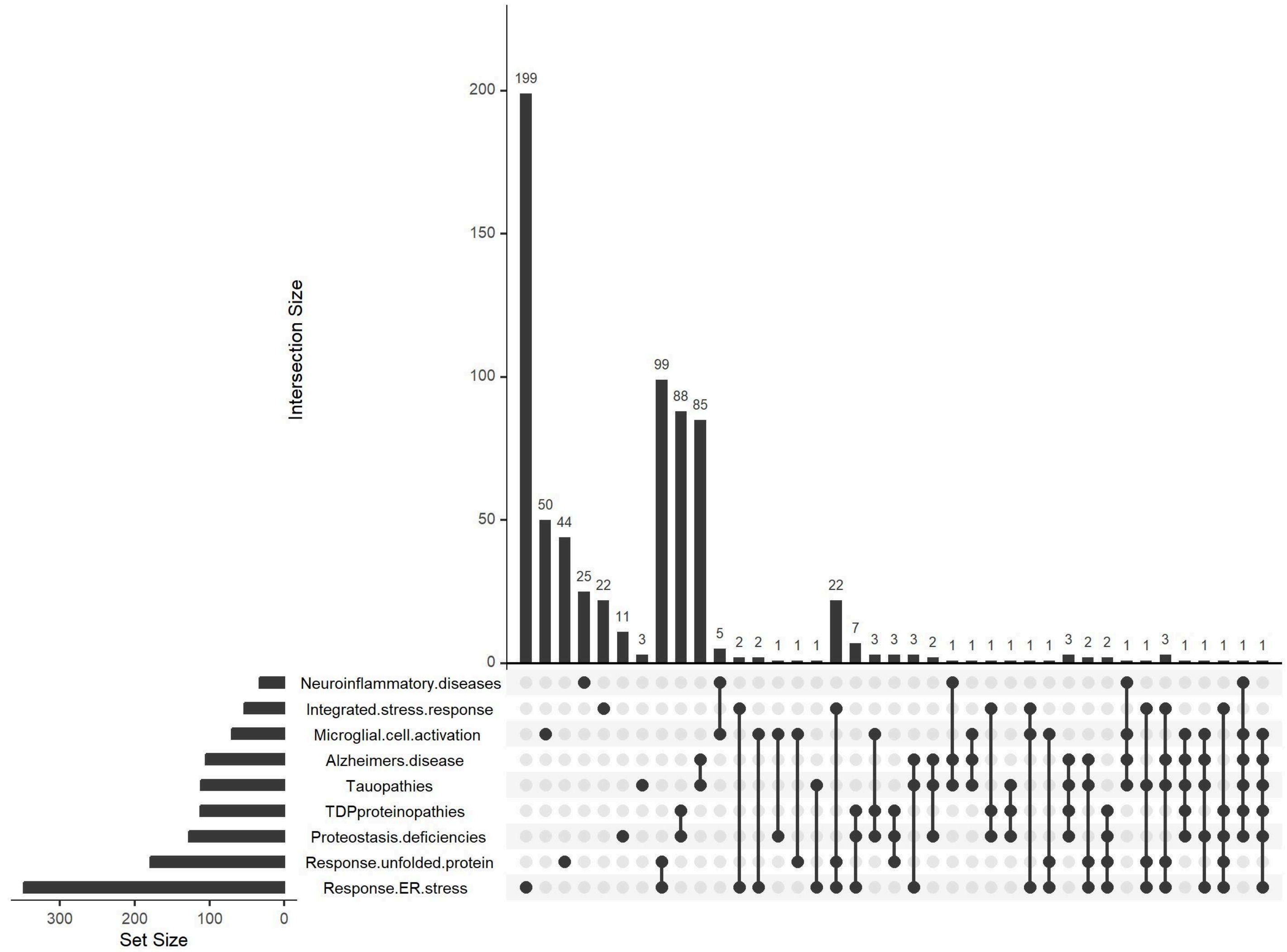


**Supplemental Figure 2.** Upset plot of the overlap in gene signatures of dementia related pathways.

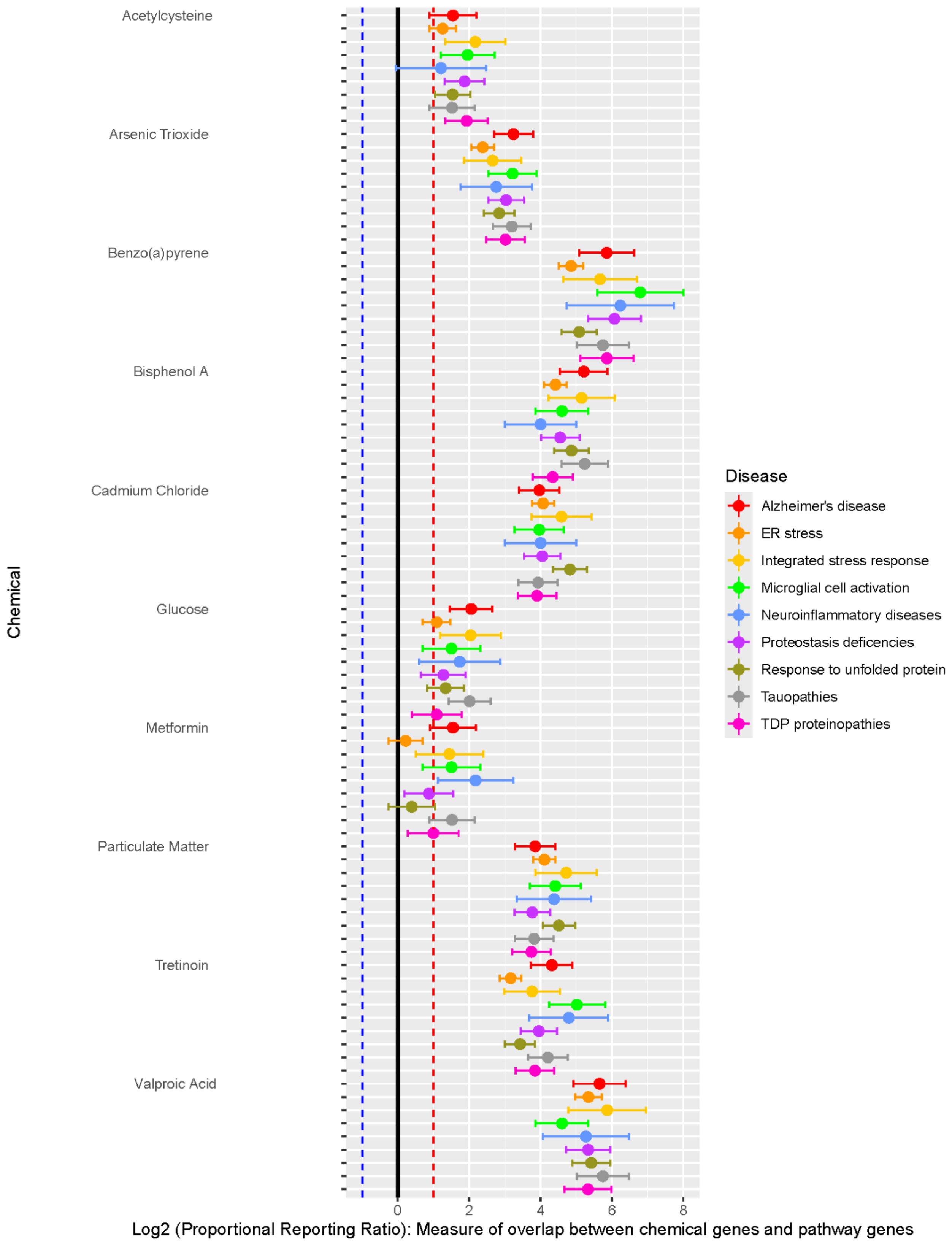


**Supplemental Figure 3.** Forest plot of Proportional Reporting Ratio (PRR) for top 10 chemicals for the nine ADRD pathways tested. Points are colored by disease list tested. Black dashed horizontal line represents PRR of zero (no enrichment or depletion), Blue dashed horizontal line represents PRR less than or equal to -1 (depletion), and the red dashed horizontal line represents PRR greater than 1 (enrichment).

**Supplementary Table 1.** List of genes for each chemical. Date of download from the Comparative Toxicogenomics Database: August 6, 2025.
CTD_supplemental_table1_11.5.25.xlsx

**Supplementary Table 2.** List of genes for each disease pathway. Date of download from the Comparative Toxicogenomics Database: August 6, 2025
CTD_supplemental_table2_10.7.25.xlsx

**Supplementary Table 3.** Results of each enrichment test for pairwise combinations of chemicals and disease pathways.
Supplemental_Table3_11.6.25.xlsx

**Supplemental Table 4.** Table showing the 15 chemicals meeting FDR<$1x10^{-6}$ threshold for all 9 disease lists, and the Proportional Reporting Ratio (PRR) for each pathway.

| Chemical name | Alzheimer's disease | ER stress | Integrated stress response | Microglial cell activation | Neuroinflam matory diseases | Proteostasis deficiencies | Response to unfolded protein | Tauopathies | TDP proteinopathies |
|---|---|---|---|---|---|---|---|---|---|
| | PRR | PRR | PRR | PRR | PRR | PRR | PRR | PRR | PRR |
| 2-(2-amino-3-methoxyphenyl)-4H-1-benzopyran-4-one | 2.9 | 0.9 | 1.6 | 2.6 | 2.8 | 1.7 | 0.7 | 2.7 | 1.9 |
| Benzo(a)pyrene | 57.9 | 28.9 | 50.9 | 111 | 75.4 | 67.3 | 33.9 | 53.7 | 58.1 |
| Capsaicin | 2.2 | 1.1 | 2.1 | 1.3 | 2.3 | 1.2 | 1 | 2 | 1.4 |
| Ethanol | 3.3 | 1.7 | 3.7 | 2.2 | 5.9 | 2.3 | 1.9 | 3.2 | 2.1 |
| Glucose | 4.2 | 2.1 | 4.1 | 2.8 | 3.3 | 2.4 | 2.5 | 4 | 2.1 |
| Glutathione | 1.6 | 0.8 | 1.8 | 1.7 | 2.3 | 1.5 | 0.7 | 1.6 | 1.6 |
| Lipopolysaccharides | 4.8 | 2.7 | 5.8 | 11 | 12.5 | 5.5 | 3.1 | 4.6 | 5.6 |
| Metformin | 2.9 | 1.2 | 2.7 | 2.8 | 4.5 | 1.8 | 1.3 | 2.9 | 2 |
| Paraquat | 2.4 | 2.1 | 3.4 | 3.9 | 5.2 | 3.2 | 2.8 | 2.6 | 3.5 |
| Particulate Matter | 14.4 | 17.2 | 26.4 | 21.3 | 20.8 | 13.6 | 22.8 | 14.2 | 13.4 |
| Quercetin | 6.4 | 4.7 | 7.4 | 7.4 | 9.8 | 7.9 | 6.4 | 6.3 | 7.5 |
| Rosiglitazone | 4.0 | 1.4 | 2.1 | 2.4 | 3.9 | 2.3 | 1.5 | 3.7 | 2.1 |
| SB 203580 | 2.3 | 1.1 | 2.7 | 1.7 | 2.3 | 1.9 | 1 | 2.1 | 2 |
| Tetradecanoylphorbol Acetate | 1.9 | 1.1 | 2.1 | 5.1 | 6.8 | 2.5 | 1.1 | 1.8 | 2.8 |
| Tretinoin | 19.9 | 9 | 13.6 | 32.4 | 27.7 | 15.5 | 10.8 | 18.5 | 14.4 |

**Supplemental Table 5.** Table showing the number of chemicals enriched and depleted (FDR<$1x10^{-6}$) by chemical category and disease list, out of the total 1,008 chemicals tested.

| Category | Total chemicals | Chemicals associated with any pathway | Alzheimer's disease | | ER stress | | Integrated stress response | | Microglial cell activation | | Neuroinflammatory diseases | | Proteostasis deficencies | | Response to unfolded protein | | Tauopathies | | TDP proteinopathies | |
|---|---|---|---|---|---|---|---|---|---|---|---|---|---|---|---|---|---|---|---|---|
| | | | Depleted | Enriched | Depleted | Enriched | Depleted | Enriched | Depleted | Enriched | Depleted | Enriched | Depleted | Enriched | Depleted | Enriched | Depleted | Enriched | Depleted | Enriched |
| Overall | 1,008 | 742 | 214 | 295 | 392 | 200 | 0 | 193 | 41 | 172 | 0 | 75 | 183 | 231 | 174 | 230 | 206 | 305 | 155 | 231 |
| Biological Factors | 48 | 38 | 11 | 16 | 21 | 7 | 0 | 14 | 2 | 14 | 0 | 9 | 11 | 9 | 8 | 9 | 12 | 16 | 8 | 10 |
| Carboxylic acids | 30 | 21 | 5 | 12 | 11 | 5 | 0 | 5 | 0 | 2 | 0 | 2 | 4 | 8 | 5 | 10 | 5 | 12 | 4 | 7 |
| Elements and Minerals | 14 | 10 | 2 | 6 | 6 | 3 | 0 | 3 | 1 | 6 | 0 | 1 | 2 | 5 | 3 | 5 | 2 | 6 | 1 | 6 |
| Environmental Pollutants and Complex Mixtures | 24 | 16 | 2 | 10 | 5 | 10 | 0 | 7 | 1 | 8 | 0 | 2 | 2 | 11 | 3 | 11 | 2 | 10 | 2 | 10 |
| Fluorine, Chlorine or Bromine compounds | 9 | 6 | 5 | 1 | 3 | 1 | 0 | 1 | 0 | 0 | 0 | 0 | 3 | 0 | 2 | 1 | 5 | 1 | 3 | 0 |
| Fluorocarbons | 9 | 4 | 1 | 2 | 0 | 3 | 0 | 0 | 0 | 2 | 0 | 1 | 0 | 2 | 0 | 3 | 1 | 2 | 0 | 2 |
| Heterocyclic Compounds | 210 | 160 | 56 | 48 | 94 | 35 | 0 | 40 | 6 | 26 | 0 | 13 | 41 | 35 | 34 | 43 | 50 | 52 | 31 | 37 |
| Hormones, Hormone Substitutes, Steroids and Hormone Antagonists | 39 | 27 | 6 | 12 | 15 | 6 | 0 | 6 | 1 | 6 | 0 | 4 | 5 | 10 | 7 | 6 | 6 | 11 | 5 | 11 |
| Hydrocarbons (Chlorinated, Acyclic, Aromatic, Halogenated, Cyclic, Other) | 37 | 30 | 8 | 12 | 18 | 6 | 0 | 6 | 2 | 5 | 0 | 1 | 8 | 8 | 8 | 7 | 8 | 12 | 7 | 9 |
| Metals and Metal Compounds | 88 | 74 | 15 | 36 | 34 | 26 | 0 | 22 | 3 | 22 | 0 | 2 | 17 | 31 | 19 | 28 | 13 | 36 | 16 | 27 |

| | | | | | | | | | | | | | | | | | | | | |
|---|---|---|---|---|---|---|---|---|---|---|---|---|---|---|---|---|---|---|---|---|
| Nucleic Acids, Nucleotides, Nucleosides, Amino Acids, Peptides, and Proteins | 37 | 28 | 7 | 10 | 14 | 7 | 0 | 8 | 0 | 8 | 0 | 4 | 5 | 9 | 5 | 7 | 6 | 10 | 5 | 9 |
| Organophosphorus compounds | 16 | 13 | 4 | 6 | 5 | 3 | 0 | 1 | 0 | 2 | 0 | 0 | 3 | 4 | 3 | 4 | 4 | 6 | 3 | 2 |
| Other (Selenium Compounds, Onium Compounds, Gases, Free Radicals, Coordination Complexes, Noxae, Boron Compounds) | 19 | 14 | 2 | 7 | 9 | 4 | 0 | 8 | 0 | 4 | 0 | 2 | 5 | 5 | 1 | 7 | 2 | 8 | 3 | 5 |
| Other Nitrogen-Containing Compounds (Amides, Amidines, Amines, Azo, Hydrazines, Isocyanates, Triazines, Nitriles, Nitro and Nitroso Compounds) | 51 | 39 | 13 | 11 | 21 | 7 | 0 | 9 | 3 | 5 | 0 | 5 | 12 | 5 | 13 | 8 | 14 | 12 | 10 | 6 |
| Other Organic Chemicals (Ethers, Alcohols, Aldehydes, Ketones, Lactones, Quinones, Carbohydrates, Lipids, Organosilicon | 106 | 69 | 27 | 21 | 43 | 14 | 0 | 17 | 8 | 17 | 0 | 7 | 20 | 21 | 22 | 14 | 26 | 23 | 16 | 20 |
| Personal Care, Consumer Products, Therapeutic Agents, Pharmacologic Actions and Specialty Use Chemicals | 48 | 41 | 10 | 20 | 16 | 15 | 0 | 12 | 6 | 13 | 0 | 6 | 11 | 15 | 9 | 14 | 11 | 20 | 10 | 15 |
| Pesticides, Pesticide Synergists and Cholinesterase Inhibitors | 16 | 11 | 0 | 6 | 5 | 4 | 0 | 3 | 0 | 3 | 0 | 2 | 1 | 6 | 3 | 5 | 0 | 6 | 2 | 5 |

| | | | | | | | | | | | | | | | | | | | | |
|---|---|---|---|---|---|---|---|---|---|---|---|---|---|---|---|---|---|---|---|---|
| Phthalic Acids and Plasticizers | 11 | 8 | 4 | 3 | 3 | 2 | 0 | 0 | 0 | 1 | 0 | 0 | 1 | 2 | 1 | 2 | 4 | 3 | 1 | 2 |
| Polychlorinated Biphenyls and Flame Retardants | 21 | 18 | 4 | 7 | 9 | 5 | 0 | 3 | 0 | 3 | 0 | 1 | 2 | 5 | 5 | 7 | 3 | 8 | 2 | 4 |
| Polycyclic Aromatic Hydrocarbons | 46 | 29 | 10 | 13 | 16 | 7 | 0 | 9 | 3 | 7 | 0 | 5 | 6 | 8 | 5 | 8 | 10 | 13 | 5 | 8 |
| Phytoestrogens, Alkaloids, and Terpenes | 69 | 44 | 11 | 21 | 27 | 13 | 0 | 12 | 1 | 10 | 0 | 4 | 13 | 15 | 9 | 13 | 11 | 22 | 13 | 16 |
| Sulfur compounds | 16 | 15 | 6 | 2 | 7 | 4 | 0 | 1 | 0 | 4 | 0 | 3 | 5 | 3 | 5 | 4 | 6 | 2 | 2 | 6 |
| Vitamins and Dietary Components | 10 | 8 | 0 | 7 | 1 | 6 | 0 | 3 | 2 | 1 | 0 | 0 | 1 | 6 | 0 | 6 | 0 | 7 | 1 | 6 |
| Volatile Organic Compounds | 10 | 6 | 2 | 1 | 3 | 2 | 0 | 0 | 0 | 2 | 0 | 0 | 1 | 2 | 0 | 2 | 2 | 1 | 1 | 2 |
| Other | 24 | 13 | 3 | 5 | 6 | 5 | 0 | 3 | 2 | 1 | 0 | 1 | 4 | 6 | 4 | 6 | 3 | 6 | 4 | 6 |

**Supplemental Table 6.** Table showing number (%) for all chemicals in the analytic sample (N=1,008) that impact specific dementia hallmark genes.

| Dementia hallmark genes | Number of chemicals | Chemical names |
|---|---|---|
| APOE | 109 (11%) | 1-Methyl-4-phenylpyridinium<br>2,3,5-trichloro-6-phenyl-(1,4)benzoquinone<br>2,5,2',5'-tetrachlorobiphenyl<br>3,4,5,3',4'-pentachlorobiphenyl<br>3-nitrobenzanthrone<br>4-(4-((5-(4,5-dimethyl-2-nitrophenyl)-2-furanyl)methylene)-4,5-dihydro-3-methyl-5-oxo-1H-pyrazol-1-yl)benzoic acid<br>4-(5-benzo(1,3)dioxol-5-yl-4-pyridin-2-yl-1H-imidazol-2-yl)benzamide<br>4-aminophenylarsenoxide<br>Acetaminophen<br>Acrolein<br>Aerosols<br>Aflatoxin B1<br>Air Pollutants<br>Arachidonic Acid<br>aristolochic acid I<br>Arsenic<br>Arsenic Trioxide<br>Aspirin<br>Atorvastatin<br>Benzo(a)pyrene<br>Bezafibrate<br>bisphenol A<br>bisphenol AF<br>bisphenol B<br>bisphenol F<br>bisphenol S<br>Cadmium<br>Caffeine<br>cerous chloride<br>chloropicrin<br>Chlorpyrifos<br>Cisplatin<br>Cobalt<br>Copper<br>corosolic acid<br>Cyclosporine<br>DDT<br>deoxynivalenol<br>Dexamethasone |

| | | |
|---|---|---|
| | | Dichlorodiphenyl Dichloroethylene<br>Diethylstilbestrol<br>Dihydrotestosterone<br>dorsomorphin<br>entinostat<br>Estradiol<br>ferrous chloride<br>Fluorides<br>Fluvastatin<br>Folic Acid<br>ginger extract<br>Glucose<br>GW 3965<br>GW 4064<br>GW 7647<br>ICG 001<br>Iron<br>Isotretinoin<br>Ivermectin<br>jinfukang<br>Lead<br>licochalcone B<br>Mercury<br>methylmercuric chloride<br>N-(2-(4-bromocinnamylamino)ethyl)-5-isoquinolinesulfonamide<br>Nickel<br>nickel sulfate<br>nivalenol<br>obeticholic acid<br>Oils, Volatile<br>Okadaic Acid<br>Oleic Acid<br>palbociclib<br>paricalcitol<br>Particulate Matter<br>pentabromodiphenyl ether<br>pentanal<br>perfluorooctane sulfonic acid<br>perfluorooctanoic acid<br>Phosphorus<br>Plant Extracts<br>Polychlorinated Biphenyls<br>polyhexamethyleneguanidine<br>Polyphenols<br>potassium perchlorate<br>propionaldehyde<br>Quercetin<br>Resveratrol<br>Rosiglitazone |

| | | |
|---|---|---|
| | | Sarin<br>Selenium<br>Simvastatin<br>Smoke<br>sodium arsenite<br>Sunitinib<br>T0901317<br>Tamoxifen<br>tert-Butylhydroperoxide<br>testosterone enanthate<br>Tetrachlorodibenzodioxin<br>Tobacco Smoke Pollution<br>tobacco tar<br>Tretinoin<br>Troglitazone<br>tungsten carbide<br>Valproic Acid<br>Valsartan<br>Warfarin<br>Zearalenone<br>Zinc |
| APP | 163 | 2-(2-amino-3-methoxyphenyl)-4H-1-benzopyran-4-one<br>2,2'-methylenebis(4-methyl-6-tert-butylphenol)<br>2-(4-morpholinyl)-8-phenyl-4H-1-benzopyran-4-one<br>27-hydroxycholesterol<br>3,3',4,5'-tetrahydroxystilbene<br>3-methyladenine<br>4-((3-bromophenyl)amino)-6,7-dimethoxyquinazoline<br>4-(5-benzo(1,3)dioxol-5-yl-4-pyridin-2-yl-1H-imidazol-2-yl)benzamide<br>4-hydroxy-2-nonenal<br>Acetaminophen<br>Acetylcysteine<br>Acrolein<br>Adenine<br>Adenosine Triphosphate<br>AICA ribonucleotide<br>Air Pollutants<br>Alitretinoin<br>alpha-pinene<br>alpha-Tocopherol<br>Aluminum<br>Arachidonic Acid<br>aristolochic acid I<br>Arsenic<br>arsenic trichloride<br>Arsenic Trioxide<br>Asbestos, Crocidolite<br>Ascorbic Acid |

| | | |
|---|---|---|
| | | Benzene<br>Benzo(a)pyrene<br>beta-lapachone<br>bisphenol A<br>bisphenol F<br>Cacodylic Acid<br>Cadmium<br>Cadmium Chloride<br>Cannabidiol<br>chloropicrin<br>Clioquinol<br>Clozapine<br>Cobalt<br>cobaltous chloride<br>Cocaine<br>Copper<br>corosolic acid<br>Coumestrol<br>Curcumin<br>Cyclosporine<br>DDT<br>decabromobiphenyl ether<br>Deferoxamine<br>Deoxyglucose<br>Dichlorodiphenyl Dichloroethylene<br>Dinoprostone<br>Disulfiram<br>dorsomorphin<br>Doxorubicin<br>Dronabinol<br>enzalutamide<br>epigallocatechin gallate<br>Estradiol<br>Fenofibrate<br>Folic Acid<br>Fonofos<br>Fulvestrant<br>Gallic Acid<br>Glucose<br>Glyphosate<br>Go 6976<br>gossypol acetic acid<br>GW 501516<br>Heparin<br>hexabromocyclododecane<br>Hydrogen Peroxide<br>Ibuprofen<br>ICG 001<br>Indomethacin |

| | | |
|---|---|---|
| | | Iron<br>Isoniazid<br>Ivermectin<br>kojic acid<br>lactacystin<br>Lactic Acid<br>LDN 193189<br>Lead<br>lead acetate<br>Lipopolysaccharides<br>Lithium Chloride<br>Lovastatin<br>Manganese<br>Melatonin<br>Menthol<br>Mercuric Chloride<br>Mercury<br>methacrylaldehyde<br>Methamphetamine<br>Methotrexate<br>Mitomycin<br>morin<br>moringin<br>MRK 003<br>Nanotubes, Carbon<br>Nickel<br>nickel acetate<br>Nicotine<br>ochratoxin A<br>Oils, Volatile<br>Orlistat<br>Ozone<br>Palmitic Acid<br>Paraquat<br>Parathion<br>Particulate Matter<br>Penicillamine<br>perfluorobutanesulfonic acid<br>perfluorooctane sulfonic acid<br>perfluorooctanoic acid<br>PF-06840003<br>Pioglitazone<br>Plant Extracts<br>potassium bromate<br>pyrazolanthrone<br>Quercetin<br>Raloxifene Hydrochloride<br>Reactive Oxygen Species<br>Resveratrol |

| | | |
|---|---|---|
| | | Ribonucleotides<br>Rosiglitazone<br>Rotenone<br>Sarin<br>SB 203580<br>Selenium<br>Serotonin<br>Sevoflurane<br>Silicon Dioxide<br>Simvastatin<br>Sirolimus<br>sodium arsenite<br>Staurosporine<br>Sulindac<br>sulindac sulfide<br>terbufos<br>tert-Butylhydroperoxide<br>testosterone enanthate<br>tetrabromobisphenol A<br>tetrathiomolybdate<br>Thiosemicarbazones<br>Tobacco Smoke Pollution<br>Tretinoin<br>Triiodothyronine<br>triphenyl phosphate<br>tris(2-butoxyethyl) phosphate<br>Troglitazone<br>Tunicamycin<br>U 0126<br>Uranium<br>uranyl acetate<br>Urethane<br>Ursolic Acid<br>Valproic Acid<br>Vehicle Emissions<br>Vitamin E<br>Volatile Organic Compounds<br>Zinc |
| PSEN1 | 52 | 4-(5-benzo(1,3)dioxol-5-yl-4-pyridin-2-yl-1H-imidazol-2-yl)benzamide<br>7,8-Dihydro-7,8-dihydroxybenzo(a)pyrene 9,10-oxide<br>Acetaminophen<br>Air Pollutants, Occupational<br>Amiodarone<br>Antirheumatic Agents<br>Atrazine<br>Benzo(a)pyrene<br>benzyloxycarbonylleucyl-leucyl-leucine aldehyde<br>bisphenol A |

| | | |
|---|---|---|
| | | Bortezomib<br>Cadmium Chloride<br>Cannabidiol<br>CGP 52608<br>Copper<br>Copper Sulfate<br>Cycloheximide<br>DDT<br>deguelin<br>deoxynivalenol<br>Dronabinol<br>enzalutamide<br>Fenofibrate<br>FR900359<br>Go 6976<br>(+)-JQ1 compound<br>Lead<br>Lipopolysaccharides<br>Menthol<br>Methotrexate<br>Methyl Methanesulfonate<br>Nickel<br>Oxaliplatin<br>Paraquat<br>Particulate Matter<br>Plant Extracts<br>Resveratrol<br>Sarin<br>SB 203580<br>sodium arsenite<br>Sodium Selenite<br>sulindac sulfide<br>Tobacco Smoke Pollution<br>Tretinoin<br>trichostatin A<br>triphenyl phosphate<br>tris(1,3-dichloro-2-propyl)phosphate<br>Tunicamycin<br>Urethane<br>Valproic Acid<br>Vehicle Emissions<br>Zinc |
